# Maternal parity affects Day 8 embryo gene expression in old mares

**DOI:** 10.1101/2021.12.01.470709

**Authors:** Emilie Derisoud, Luc Jouneau, Clothilde Gourtay, Anne Margat, Catherine Archilla, Yan Jaszczyszyn, Rachel Legendre, Nathalie Daniel, Nathalie Peynot, Michèle Dahirel, Laurie Briot, Frédéric De Geoffroy, Véronique Duranthon, Pascale Chavatte-Palmer

**Affiliations:** Université Paris-Saclay, UVSQ, INRAE, BREED, Jouy-en-Josas, France; Ecole Nationale Vétérinaire d’Alfort, BREED, Maisons-Alfort, France; IFCE, Plateau technique du Pin, Exmes, France; Institute for Integrative Biology of the Cell (I2BC), UMR 9198 CNRS, CEA, Paris-Sud University, Gif-sur-Yvette, France; Institut Pasteur—Bioinformatics and Biostatistics Hub—Department of Computational Biology, USR 3756 IP CNRS, Paris, France

**Keywords:** Blastocyst, RNA sequencing, horse, equine

## Abstract

As sport career is a priority in most of equine breeds, mares are frequently bred for the first time at an advanced age. Both age and first gestation were shown to have a deleterious effect on reproduction outcomes, respectively on fertility and offspring weight but the effect mare’s parity in older mares on embryo quality has never been considered. The aim of this project was to determine the effect of old mare’s nulliparity on gene expression in embryos. Day 8 post ovulation embryos were collected from old (10-16 years old) nulliparous (ON, N=5) or multiparous (OM, N=6) non-nursing Saddlebred mares, inseminated with the semen of one stallion. Pure (TE_part) or inner cell mass enriched (ICMandTE) trophoblast were obtained by embryo bisection and paired end, non-oriented RNA sequencing (Illumina, NextSeq500) was performed on each hemi-embryo. To discriminate gene expression in the ICM from that in the TE, deconvolution (DeMixT R package) was used on the ICMandTE dataset. Differential expression was analyzed (DESeq2) with embryo sex and diameter as cofactors using a false discovery rate <0.05 cutoff. Although the expression of only a few genes was altered by mare’s nulliparity (33 in ICM and 23 in TE), those genes were related to nutrient exchanges and responses to environment signaling, both in ICM and TE, suggesting that the developing environment from these mares are not optimal for embryo growth. In conclusion, being nulliparous and old does not seem to be the perfect match for embryonic development in mares.

**Summary sentence:** Mare’s parity in old mares impacts the expression of genes related to development and molecule exchanges in ICM and TE of blastocysts suggesting an adaptation to an altered environment.

## Introduction

In the equine industry, mares are bred until an advanced age for economic and sentimental reasons. Depending on the breed, mares older than 10 years old represent between 37 and 63% of the broodmares [1-3]. In addition, 4% of Thoroughbred mares in the UK are older than 18 years old at the time of covering [1].

Several reproductive parameters are affected by age and mares older than 10 years old can be considered as already old for reproduction as their fertility has already started to progressively decline [for review 4]. Oocytes are particularly affected by maternal age with alterations of spindle stability [5-7], altered gene expression [8,9], and altered metabolism [8,10–12] being reported, all suggesting that oocyte developmental potential is reduced in old mares. The resulting embryos were smaller at the same developmental age in most studies [13-23] with altered gene expression [23] and metabolism [11,12], suggesting impaired development that has mainly been related to the oocyte quality. The reproductive tract, however, is also affected by maternal age. Indeed, more oviductal masses [24], uterine morphological degenerations such as cysts [25-30] and fibrosis [31-33] as well as more endometritis [19,25,34,35] are observed in old mares.

In most of farm animals, female parity, defined as the number of pregnancies that reached a viable gestational age (stillbirth and live birth included), is highly correlated with age as to remain profitable and stay in the farm, females must produce offspring regularly. In horses, however, the sport career is prioritized and depending on discipline and breed, can last up to 15 years or even more, as in warmblood dressage and show jumping. In Finn horse and Standardbred, nulliparous mares represented, respectively, 20.5% and 15.5% of mares that were bred in Finland [2]. To the authors’ knowledge, the effect of nulliparity/primiparity on oocyte and embryo quality as well as fetal development has not been explored. In any case, the reproductive tract is affected by mare parity. Indeed, the ventral position of the uterus [36] and the number of vascular degenerations in the endometrium [37] have been positively correlated with the number of foals. Furthermore, cervical dilatation is poorer in nulliparous *vs* multiparous mares [38] and it was suggested that uterine clearance was impeded. Primiparity, however, does not affect the prevalence of endometritis [34].

Both age and parity thus affect mare reproductive efficiency but the cumulative effect of nulliparity and aging has not been explored.

The aim of this study was to determine the effect of maternal nulliparity in old mares on embryo gene expression at the blastocyst stage. Old (>10 years) nulliparous and multiparous mares were inseminated with the same stallion semen. Blastocysts were collected and bisected to separate the pure trophoblast (TE_part) from the inner cell mass enriched hemi-embryo (ICMandTE). Gene expression was analyzed by RNA-seq in each compartment.

## Materials and methods

### Ethics

The experiment was performed at the experimental farm of IFCE (“Institut Français du Cheval et de l’Equitation – La jumenterie du Pin” research agreement D61-157-2 valid until November 2023). The protocol was approved by the local animal care and use committee and by the regional ethical committee (“Comité d’Éthique Normand en Matière d’Expérimentation Animale”, approved under N° CEEA - 54 in the National Registry of French Ethical Committees for animal experimentation) under protocol number APAFIS#20857-2019051709319621 v3. All experiments were performed in accordance with the European Union Directive 2010/63EU.

### Embryo collection

Twenty-five multiparous mares (mostly Selle Français breed with some French Anglo-Arabian, Standardbred and Saddlebred) aged from 10 to 16 years old were included in this study. Multiparous mares were defined as dams that had already foaled at least once while nulliparous mares were defined as mares that have never foaled before the experiment. During the experimental protocol, mares were managed in two herds, independent of mare group, in natural pastures 24h/day with free access to water. The experiments took place from July 8^th^ to August 13^th^, 2019. All mares remained healthy during this period.

Mares were allocated to one of 2 groups according to their parity: nulliparous (ON, n = 11) and multiparous mares (OM, n = 14). During the experimentation, mare’s withers’ height and weight were measured. Characteristics of all mares and mares that produced an embryo are detailed in Table 1.

**Table 1:** Mares’ characteristics at embryo collection time.

| Characteristics | Nulliparous (ON) |  | Multiparous (OM) |  |
| --- | --- | --- | --- | --- |
|  | All mares<br>(n = 11) | Embryo donors<br>(n = 5) | All mares<br>(n = 14) | Embryo donors<br>(n = 6) |
| Breed | SF n = 9; ST n = 1;<br>SB n = 1 | SF n = 4; ST n = 1 | SF n = 7; AA n = 4;<br>ST n = 2 | SF n = 5; AA n = 1 |
| Age (in years) | 12.7 ± 2.0 | 13.6 ± 2.2 | 13.4 ± 2.1 | 13.5 ± 2.4 |
| Parity (number of foalings) | 0 | 0 | 1.7 ± 1.3 | 1.8 ± 1.6 |
| Weight (in kg) | 663.3 ± 49.1 | 644.8 ± 53.3 | 641.0 ± 76.1 | 658.0 ± 81.7 |
| Withers' height (in cm) | 168.2 ± 5.8 | 165.8 ± 7.6 | 163.6 ± 4.1 | 164.0 ± 5.1 |
| Progesterone concentration (in ng/ml) | - | 12.7 ± 5.8 | - | 14.5 ± 7.3 |

All mares were at least 10 years old.
AA: Anglo Arab or Anglo-Arabian type; SF: Selle Français section A or B; ST: Standardbred; SB:
Saddlebred. Age, parity, weight, and height are presented as mean ± SD

The mares’ estrous period was monitored routinely in the morning by ultrasound with a 50-60Hz trans-rectal transducer. During estrus, ovulation was induced with a single injection of human chorionic gonadotropin (i.v.; 1500IU; Chorulon® 5000; MSD Santé animale, France) as soon as one ovarian follicle > 35mm in diameter was observed, together with marked uterine edema. Ovulation usually takes place within 48h, with > 80% occurring 25 to 48h after injection [39,40]. At the same time and one day later, mares were inseminated once with fresh or refrigerated semen containing at least 1 billion motile spermatozoa from a single fertile stallion. Ovulation was checked every 12-24 hours by ultrasonography. If no embryo was recovered, the procedure could be repeated once more.

Embryos were collected by non-surgical uterine lavage using prewarmed (37°C) lactated Ringer’s solution (B.Braun, France) and EZ-Way Filter (IMV Technologies, France) on the morning, 8 days post ovulation. At Day 14 post ovulation, a pregnancy diagnosis was performed for each mare and they were treated with luprotiol, an analogue of prostaglandin F2α (i.m; 7.5 mg; Prosolvin, Virbac, France).

When an embryo was collected, a blood sampling was performed at the same time on heparin tube. Plasma was recovered after centrifugation (3500 rpm, 10min at 4°C). Progesterone was measured in plasma using ELISA assay as previously described [41,42].

### Embryo bisection and RNA extraction

Using a binocular magnifying glass, collected embryos were immediately photographed with a size standard to subsequently determine embryo diameter using ImageJ® software (version 1.52a; National Institutes of Health, Bethesda, MD, USA). Then embryos were washed 4 times in commercially available Embryo holding medium (IMV Technologies, France) at 34°C and bisected with a microscalpel under binocular magnifying glass to obtain a trophoblast (TE_part) and an inner cell mass enriched (ICMandTE) hemi-embryo. At this stage, the TE_part is composed of trophectoderm and endoderm whereas the ICM is composed of epiblast cells. Directly after bisection, RNA extraction of each hemi-embryo was started in extraction buffer (PicoPure RNA isolation kit, Applied Biosystems, France) for 30 min at 42°C prior to storage at - 80°C. RNA was extracted later on from each hemi-embryo using PicoPure RNA isolation kit (PicoPure RNA isolation kit, Applied Biosystems, France), which included a DNAse treatment, following the manufacturer’s instructions. RNA quality and quantity were assessed with the 2100 Bioanalyzer system using RNA 6000 Pico kit (Agilent Technologies, France) according to the manufacturer’s instructions.

### RNA sequencing

Five nanograms of total RNA were mixed with ERCC spike-in mix (Thermofisher Scientific, France) according to manufacturer’s recommendations. Messenger RNAs were reverse transcribed and amplified using the SMART-Seq V4 ultra low input RNA kit (Clontech, France) according to the manufacturer recommendations. Nine PCR cycles were performed for each hemi-embryo. cDNA quality was assessed on an Agilent Bioanalyzer 2100, using an Agilent High Sensitivity DNA Kit (Agilent Technologies, France). Libraries were prepared from 0.15 ng cDNA using the Nextera XT Illumina library preparation kit (Illumina, France). They were pooled in equimolar proportions and sequenced (Paired end 50-34 pb) on NextSeq500 instrument, using a NextSeq 500 High Output 75 cycles kit (Illumina, France). Demultiplexing was performed with bcl2fastq2 version 2.2.18.12 (Illumina, France) and adapters were trimmed with Cutadapt version 1.15 [43]. Only reads longer than 10pb were kept.

### RNA mapping and counting

As previously described [23], alignment was performed using STAR version 2.6 [44] on previously modified Ensembl 99 EquCab3.0 assembly and annotation. Genes were then counted with FeatureCounts [45] from Subreads package version 1.6.1.

### Availability of data and materials

The RNA sequencing data supporting the conclusions of this article are available in the GEO repository, [accession: GSE188866; https://www.ncbi.nlm.nih.gov/geo/query/acc.cgi?acc=GSE188866].

### Data analysis

All statistical analyses were performed by comparing ON to OM (set as reference group) using R version 4.0.2 [46] on Rstudio software version 1.3.1056 [47].

Embryo were sexed using *X Inactive Specific Transcript* (*XIST*) expression as previously described [23]. Seven embryos were determined as females (4 in the ON group and 3 in the OM group) while 4 were considered as males (1 in the ON group, and 3 in the OM group).

### Embryo recovery and fertility rate, embryo diameter and total RNA content analysis

Embryo recovery rates (ERR) were calculated as the number of attempts with at least one embryo collected/total number of attempts. Furthermore, fertility was calculated as the sum of embryo collections with at least one embryo and the number of positive pregnancy checks at Day 11 after a negative embryo collection on the total number of attempts. Both were analyzed using the Exact Fisher test to determine if maternal parity in old mares influenced embryo recovery and the probability of leaving an embryo in the uterus after uterine flush.

For total RNA content analyses, as embryos were bisected without strict equality for each hemi-embryo, a separate analysis of ICMandTE and TE_part RNA quantities would not have been meaningful. Thus, ICMandTE and TE_part RNA quantities were summed up. With embryo diameter, both variables were analyzed using a linear model of nlme package version 3.1-148 [48] including maternal age and embryo sex, followed by 1000 permutations using PermTest function from pgirmess package version 1.6.9 [49]. Variables were kept in models when statistically significant differences were observed. Differences were considered as significant for p < 0.05.

### Deconvolution of gene expression in ICMandTE using DeMixT

The deconvolution method has already been described in equine embryos [23]. Briefly, this method enables the estimation of the relative gene expression of TE and ICM cell types within the hemi-embryo ICMandTE which is composed of both trophoblast and inner cell mass in unknown relative proportions. After filtering out all genes with at least 3 null count values in at least one group (ON or OM) per hemi-embryo (ICMandTE or TE_part), removing genes with a null variance in TE_part and adding the value “1” to all count values in ICMandTE and TE_part datasets, deconvolution was performed using the DeMixT R package version 1.4.0 [50,51]. Output datasets were DeMixT_ICM_cells and DeMixT_TE_cells, corresponding to the deconvoluted gene expression in ICM cells and TE cells of ICMandTE, respectively.

At the end of deconvolution, a quality check was automatically performed by the DeMixT R package with the TE_part used as reference for DeMixT_TE_cells. Genes were automatically filtered out if the difference between average deconvoluted expression of reference cells in mixed samples and average expression of reference cells > 4.

Outputs of DeMixT_ICM_cells *vs* DeMixT_TE_cells, DeMixT_ICM_cells *vs* TE_part and ICMandTE *vs* TE_part were compared with Deseq2 version 1.28.1 [52] to confirm that the deconvolution was effective at separating gene expression. To check if deconvolution was efficient, as previously described [23], the expression of several genes proper to ICM and TE cells in equine embryos identified using literature search [53] was compared before and after deconvolution. Results of these analyses were represented through manually drawn Venn diagrams as well as principal component analysis graphics of individuals, using ggplot2 version 3.3.3 [54] and factoextra version 1.0.7 [55].

### Maternal parity comparison for gene expression

All genes with an average expression <10 counts in both ON and OM per hemi-embryo (ICM or TE) were filtered out on the DeMixT_ICM_cells and TE_part datasets. Differential analyses were performed with Deseq2 version 1.28.1 [52] with the OM group as reference, without independent filtering and taking into account embryo diameter and sex in the model. Genes were considered differentially expressed (DEG) for FDR < 0.05 after Benjamini-Hochberg correction (also known as false discovery rate, FDR).

Equine Ensembl IDs were converted into Human Ensembl IDs and Entrez Gene names using gorth function in gprofiler2 package version 0.1.9 [56]. Genes without Entrez Gene names using gprofiler2 were manually converted when Entrez Gene names were available, using Ensembl web search function [57]. GO molecular function and GO Biological process annotations were obtained from Uniprot website.

### Gene set enrichment analyses (GSEA)

After log transformation using RLOG function of DESeq2 version 1.28.1, gene set enrichment analyses (GSEA) were performed on expressed genes using GSEA software version 4.0.3 (Broad Institute, Inc., Massachusetts Institute of Technology, and Regents of the University of California) [58,59] to identify biological gene sets disturbed by maternal parity. Molecular Signatures Databases [60] version 7.1 (C2: KEGG: Kyoto Encyclopedia of Genes and Genomes; REACTOME, C5: BP: GO biological process) were used to identify most perturbed pathways. Pathways were considered significantly enriched for FDR< 0.05. When the normalized enrichment score (NES) was positive, the gene set was enriched in the ON group while when NES was negative, the gene set was enriched in the OM group.

## Results

### Embryo recovery rates, diameter, total RNA content and quality and progesterone concentrations

Altogether, 32 embryo collections were performed (14 in ON and 18 in OM, 8 mares being flushed twice) and 15 embryos were obtained (6 from 5 ON mares and 9 from 8 OM mares). Two mares (one in each parity group) produced twin embryos.

Positive embryo collection rate was 36% and 44% in ON and OM, respectively and did not differ between groups (p = 0.72). The embryo recovery rate was 43% and 50% in ON and OM, respectively and did not differ between groups (p = 0.30). At the Day 14 pregnancy check, embryos were found in 3 OM mares (from 2 to 4 foalings) and none was found in the ON group. Fertility, calculated combining positive embryo collections and Day 14 pregnancy diagnosis, was 36% and 61% in ON and OM, respectively, and did not differ between groups (p = 0.29).

Altogether, 7 and 11 double ovulations were observed, respectively, in ON and OM. The embryo recovery rate per ovulation at the time of embryo collection and after Day 14 pregnancy check were not different according to group (respectively, 29% and 29% in ON and 31% and 41% in OM, p = 1 and p = 0.39).

All embryos were expanded blastocysts grade I or II according to the embryo classification of McKinnon and Squires [61]. For each twin collection, one embryo was large and the other was small (766µm and 295µm; 829µm and 481µm). For both, as only one embryo per mare was required, only the largest embryo of the twins was chosen for further analysis. Altogether, all ON embryos but only 6 OM embryos out of 8 collected were RNA sequenced. The smallest OM embryo (480µm) and another one randomly chosen (907µm) of diameter were not sequenced.

In embryos selected for RNA sequencing, embryo diameter ranged from 562 µm to 1426 µm, with no effect of group on embryo diameter (p = 0.18). Female embryos, however, were significantly smaller than male embryos (in average 764 ± 223 µm and 1046 ± 287 µm, p < 0.05) without interaction between maternal parity and embryo sex. RNA yield per embryo ranged from 25.2 ng to 624 ng and was not related to parity (p = 0.43) nor embryo sex (p = 0.08).

The median RNA Integrity Number (RIN) was 9.7 (8.9 - 10 range). Between 34.6 and 54.1 million reads per sample were obtained after trimming. On average, 74.10% of the reads were mapped on the modified EquCab 3.0 using STAR and 67.07% were assigned to genes by featureCounts. Except one old multiparous mare that had a progesterone plasma concentration of 3.9 ng/ml, progesterone concentrations in plasma were > 4 ng/ml for all mares (range from 8.3 to 25.6ng/ml with an average of 13.7ng/ml) and were not affected by mares’ parity (p = 0.66).

### Deconvolution of gene expression to discriminate ICM and TE gene expression in ICMandTE hemi-embryos

After selecting genes with less than 3 non null count values in at least one group (ON or OM) per hemi-embryo (ICMandTE or TE_part), 16,803 genes were conserved for deconvolution. In addition, nine genes were removed because their variance was null in the TE_part, as DeMixT does not allow the use of genes with a null variance in the pure sample. For these genes, the mean count in ICMandTE samples was lower or equal to 10 counts. One further gene was removed during the deconvolution because the deconvolution quality for this gene was not sufficient. Therefore, at the end of the deconvolution algorithm, 16,793 genes were available for differential analysis.

Before deconvolution, 303 genes were differentially expressed (FDR < 0.05) between the ICMandTE and the TE_part (Figure 1A). After deconvolution, the comparison between DeMixT_ICM_cells and DeMixT_TE_cells yielded 7,116 differentially expressed genes while the comparison DeMixT_ICM_cells *vs* TE_part yielded 5,615 differentially expressed genes, with 5,103 in common (74%). Moreover, all but one of the initially 303 differentially expressed genes before deconvolution were also identified as differentially expressed in both post-deconvolution analyses. On the PCA graph of individuals, Axis 1 (21.8% of variance) separated well groups according to data origin. ICMandTE and TE_part were separated on axis 1 but very close before deconvolution (Figure 1B). DeMixT_TE_cells and TE_part were partly superposed, indicating that datasets before and after deconvolution have a similar global gene expression; whereas the DeMixT_ICM_cells group is clearly separated from both, indicating that the deconvolution effectively enabled the separation of gene expression in the two cell types.

**Figure 1:**
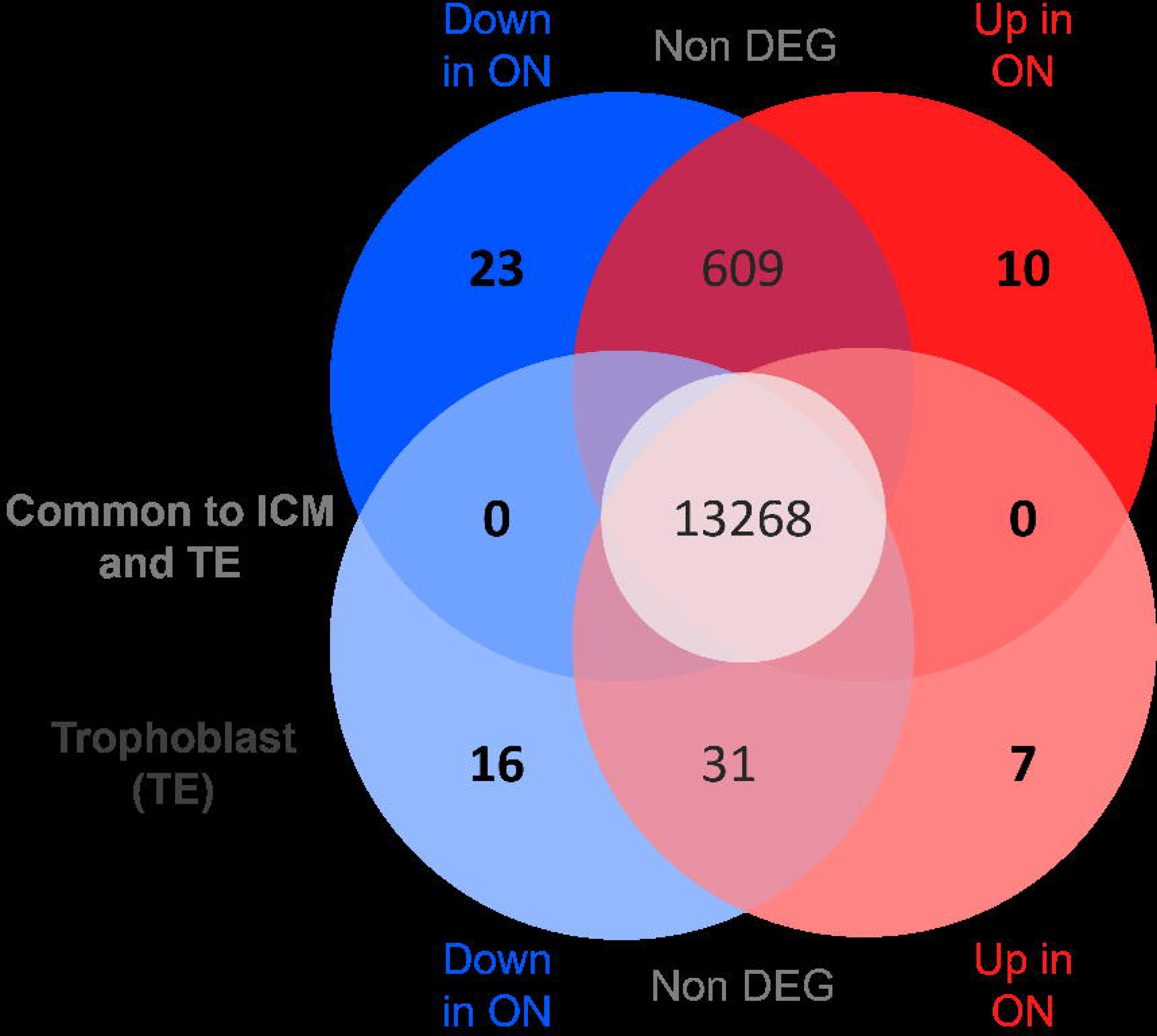
Gene expression in ICM and TE before and after deconvolution using DeMixT A) Venn diagrams of the differential gene expression in ICMandTE *vs* TE_part (before deconvolution), DeMixT_ICM_cells *vs* DeMixT_TE_cells (after deconvolution) and DeMixT_ICM_cells *vs* TE_part (gene expression of ICM after deconvolution *vs* gene expression in TE_part without deconvolution); B) Principal Component Analysis of gene expression of DeMixT_ICM_cells, DeMixT_TE_cells, ICMandTE and TE part datasets. Deconvolution was used to isolate gene expression of ICM and TE cells in ICMandTE hemi-embryos. ICMandTE: inner cell mass + trophoblast; TE_part: pure trophoblast. Here trophoblast represents trophectoderm + endoderm.

Only 5 of the 12 genes previously reported as more expressed in the ICM [53] were also identified more expressed in the ICMandTE *vs* TE_part comparison (Table 2). After deconvolution (comparison DeMixT_ICM_cells *vs* TE_part), 10 out of 12 of these genes were observed differentially expressed with 9 effectively more expressed in the ICM. The expression of *Undifferentiated Embryonic Cell Transcription Factor 1*, *UTF1*, however, was identified decreased in the DeMixT_ICM_cells, in contrast to the only published report [53]. In the TE, no gene previously identified was observed differentially expressed in the comparison ICMandTE *vs* TE_part, *i.e.*, before deconvolution. After deconvolution, the expression of 3 of the 7 reported genes were found increased in TE_part compared to DeMixT_ICM_cells.

**Table 2:**
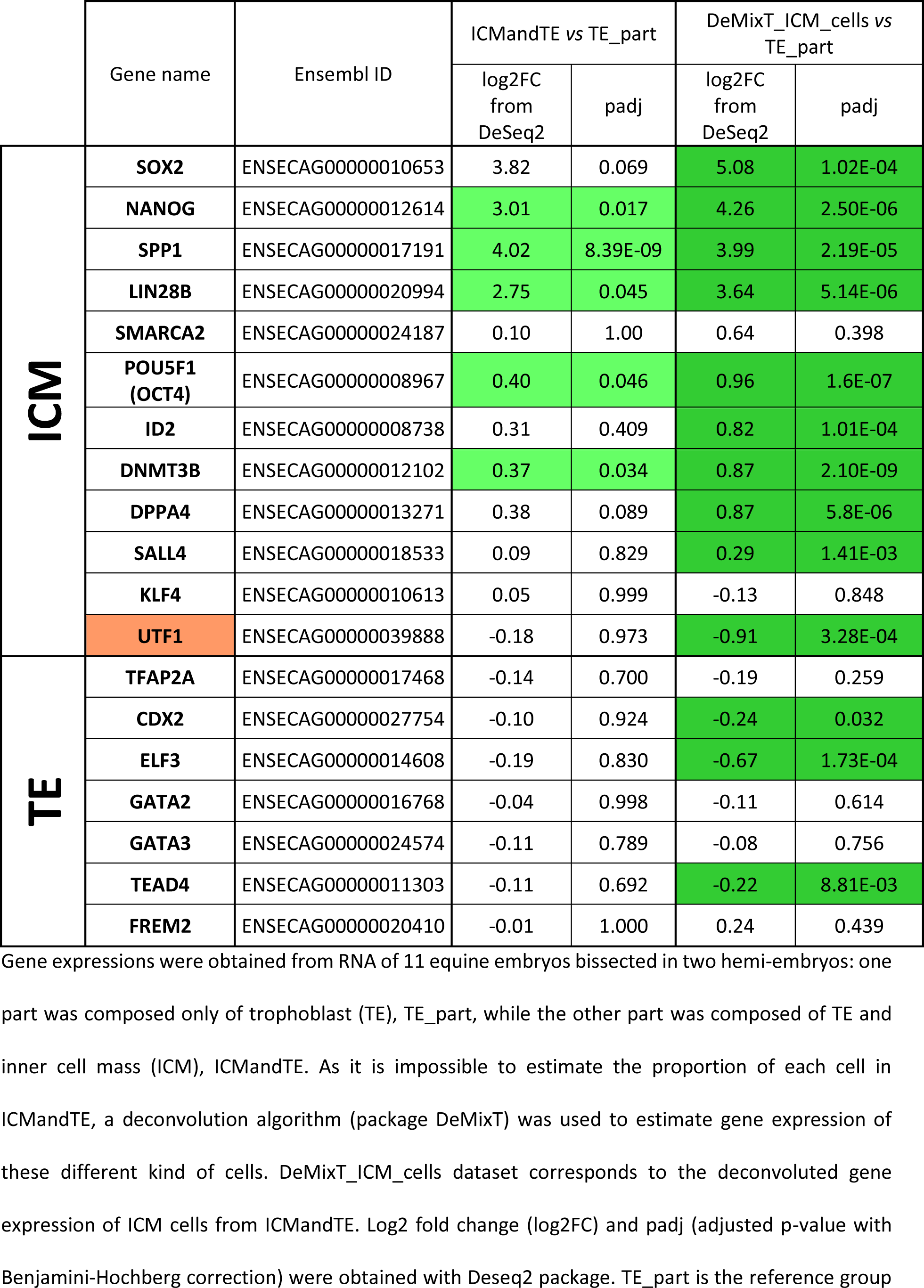
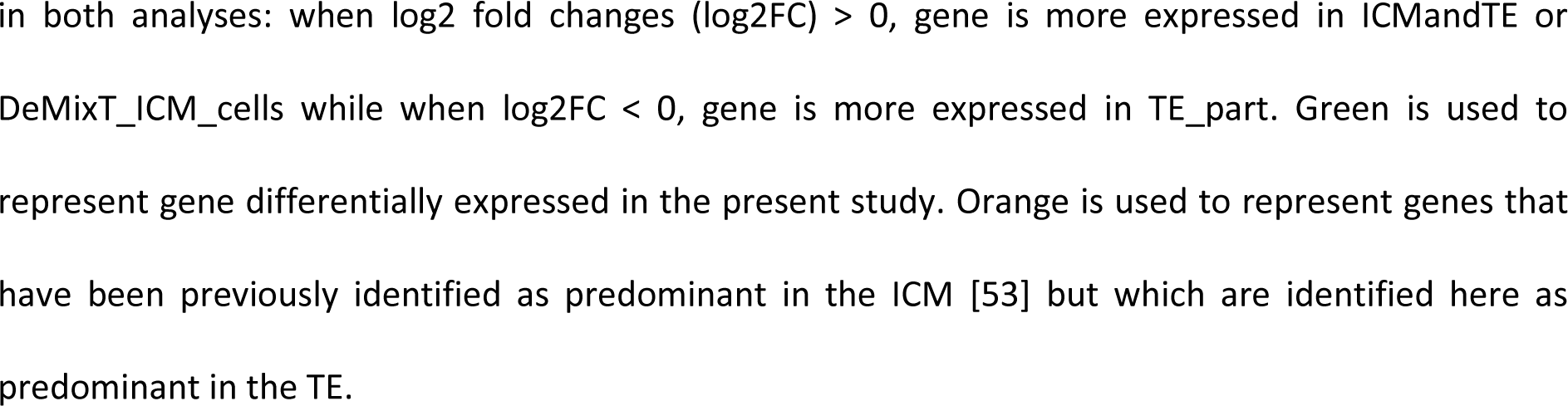
Comparison of selected genes expression before and after deconvolution

|  | Gene name | Ensembl ID | ICMandTE vs TE_part |  | DeMixT_ICM_cells vs TE_part |  |
| --- | --- | --- | --- | --- | --- | --- |
|  |  |  | log2FC from DeSeq2 | padj | log2FC from DeSeq2 | padj |
| **ICM** | **SOX2** | ENSECAG00000010653 | 3.82 | 0.069 | 5.08 | 1.02E-04 |
|  | **NANOG** | ENSECAG00000012614 | 3.01 | 0.017 | 4.26 | 2.50E-06 |
|  | **SPP1** | ENSECAG00000017191 | 4.02 | 8.39E-09 | 3.99 | 2.19E-05 |
|  | **LIN28B** | ENSECAG00000020994 | 2.75 | 0.045 | 3.64 | 5.14E-06 |
|  | **SMARCA2** | ENSECAG00000024187 | 0.10 | 1.00 | 0.64 | 0.398 |
|  | **POU5F1 (OCT4)** | ENSECAG00000008967 | 0.40 | 0.046 | 0.96 | 1.6E-07 |
|  | **ID2** | ENSECAG00000008738 | 0.31 | 0.409 | 0.82 | 1.01E-04 |
|  | **DNMT3B** | ENSECAG00000012102 | 0.37 | 0.034 | 0.87 | 2.10E-09 |
|  | **DPPA4** | ENSECAG00000013271 | 0.38 | 0.089 | 0.87 | 5.8E-06 |
|  | **SALL4** | ENSECAG00000018533 | 0.09 | 0.829 | 0.29 | 1.41E-03 |
|  | **KLF4** | ENSECAG00000010613 | 0.05 | 0.999 | -0.13 | 0.848 |
|  | **UTF1** | ENSECAG00000039888 | -0.18 | 0.973 | -0.91 | 3.28E-04 |
| **TE** | **TFAP2A** | ENSECAG00000017468 | -0.14 | 0.700 | -0.19 | 0.259 |
|  | **CDX2** | ENSECAG00000027754 | -0.10 | 0.924 | -0.24 | 0.032 |
|  | **ELF3** | ENSECAG00000014608 | -0.19 | 0.830 | -0.67 | 1.73E-04 |
|  | **GATA2** | ENSECAG00000016768 | -0.04 | 0.998 | -0.11 | 0.614 |
|  | **GATA3** | ENSECAG00000024574 | -0.11 | 0.789 | -0.08 | 0.756 |
|  | **TEAD4** | ENSECAG00000011303 | -0.11 | 0.692 | -0.22 | 8.81E-03 |
|  | **FREM2** | ENSECAG00000020410 | -0.01 | 1.000 | 0.24 | 0.439 |

Gene expressions were obtained from RNA of 11 equine embryos bisected in two hemi-embryos: one part was composed only of trophoblast (TE), TE\_part, while the other part was composed of TE and inner cell mass (ICM), ICMandTE. As it is impossible to estimate the proportion of each cell in ICMandTE, a deconvolution algorithm (package DeMixT) was used to estimate gene expression of these different kind of cells. DeMixT\_ICM\_cells dataset corresponds to the deconvoluted gene expression of ICM cells from ICMandTE. Log2 fold change (log2FC) and padj (adjusted p-value with Benjamini-Hochberg correction) were obtained with Deseq2 package. TE\_part is the reference group
in both analyses: when $\log_2$ fold changes ( $\log_2FC$ ) $> 0$ , gene is more expressed in ICMandTE or DeMixT\_ICM\_cells while when $\log_2FC < 0$ , gene is more expressed in TE\_part. Green is used to represent gene differentially expressed in the present study. Orange is used to represent genes that have been previously identified as predominant in the ICM [53] but which are identified here as predominant in the TE.

These results indicated that a better qualification of genes expressed by ICM cells was enable by the deconvolution. Thus, for further analyses, TE_part and DeMixT_ICM_cells datasets have been studied.

### Differential gene expression in deconvoluted ICM cells

After the filtering out of genes without an average expression ≥ 10 counts in at least one maternal age group/hemi-embryo, 13,910 genes were considered as expressed in the ICM cells from ON or OM embryos. Only 33 genes were differentially expressed (23 downregulated and 10 upregulated in ON) (Figure 2 and Additional file 1). Respectively, 20 and 5 genes out of the down- and upregulated genes were associated to a protein known and described in human. These 25 genes are presented in Table 3. Down regulated genes in the ICM of old nulliparous mares were involved in RNA processing and transcription, immunity, nervous system development, lipid/protein transport, lipid metabolism, chromatin remodeling, DNA repair, cell cycle, signaling and adhesion whereas up-regulated genes were related to different biological processes such as ion transport, regulation of transcription and lipid/protein/carbohydrate catabolism.

**Figure 2:**
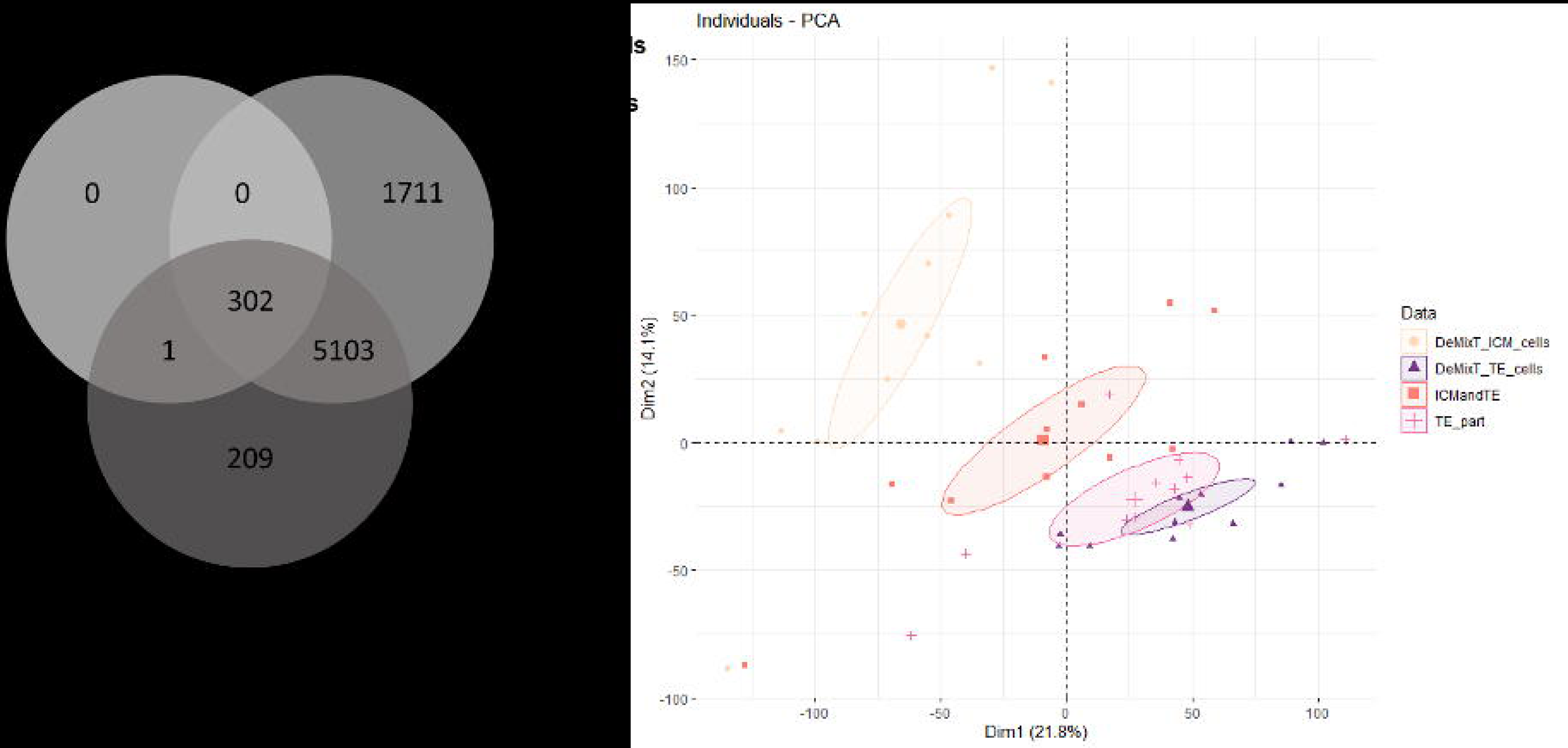
Analysis of differentially expressed genes (DEG) in embryos according to maternal parity A) representation of down- (blue) and upregulated (red) DEG in ICM (from DeMixT_ICM_cells data obtained after deconvolution of ICMandTE using DeMixT R package [50,51]) and TE (from TE_part dataset) of embryos from ON *vs* OM. DEG: Differentially Expressed Genes (FDR < 0.05); TE: Trophoblast; ICM: Inner Cell Mass; ON: Old nulliparous mares; OM: Old multiparous mares

**Table 3:** Up- and down-regulated genes coding for a protein in the inner cell mass of equine embryos according to old mare parity

| Ensembl Name | Entrez Gene ID | Description | GO Molecular function | GO Biological process | log2 Fold Change | Adjusted p-value |
| --- | --- | --- | --- | --- | --- | --- |
| ENSECAG00000002796 | GABRR1 | Gamma-aminobutyric acid type A receptor subunit rho1 | Chloride channel<br>Ion channel | Ion transport<br>Chloride transmembrane transport<br>Gamma-aminobutyric acid signaling pathway<br>Regulation of membrane potential<br>Signal transduction | 5.19 | 0.004 |
| ENSECAG00000007423 | ZKSCAN2 | zinc finger with KRAB and SCAN domains 2 | DNA-binding transcription factor activity, RNA polymerase II-specific<br>Metal ion binding | Regulation of transcription by RNA polymerase II | 4.42 | 0.008 |
| ENSECAG00000012957 | RHAG | Rh associated glycoprotein | Ammonium transmembrane transporter activity<br>Ankyrin binding<br>Leak channel activity | Ammonia transport<br>Cellular ion homeostasis<br>Bicarbonate transport<br>Carbon dioxide transport | 4.25 | 0.024 |
| ENSECAG00000017370 | PPT1 | palmitoyl-protein thioesterase 1 | Hydrolase | Nervous system development<br>Lipid catabolism process<br>Protein transport<br>Protein catabolic process | 1.97 | 0.008 |
| ENSECAG00000023194 | XYLB | xylulokinase | ATP binding<br>Kinase<br>Transferase | Carbohydrate metabolism<br>Xylose metabolism | 0.63 | 0.009 |
| ENSECAG00000036419 | RPL36 | ribosomal protein L36 | Ribonucleoprotein<br>Ribosomal protein | Translation<br>Nuclear-transcribed mRNA catabolic process, nonsense-mediated decay<br>rRNA processing | -0.36 | 0.001 |
| ENSECAG00000012792 | MIF | macrophage migration inhibitory factor | Cytokine<br>Isomerase | Immunity<br>Inflammatory response<br>Innate immunity | -0.39 | 0.001 |
| ENSECAG00000007113 | ATXN10 | ataxin 10 | Enzyme binding | Nervous system development<br>Cilium assembly<br>Neuron projection development | -0.46 | 3.78E-08 |
| ENSECAG00000020469 | VPS51 | VPS51 subunit of GARP complex |  | Lipid transport<br>Protein transport<br>Transport<br>Autophagy<br>Brain morphogenesis<br>Golgi organization | -0.56 | 0.024 |
| ENSECAG00000006539 | EXOSC7 | exosome component 7 | RNA-binding | rRNA processing | -0.65 | 0.002 |
| ENSECAG00000042897 | H2AC20 | H2A clustered histone 20 | DNA-binding | Chromatin silencing | -0.76 | 0.007 |
| ENSECAG00000020116 | RABEP2 | rabaptin, RAB GTPase binding effector protein 2 | Growth factor activity<br>GTPase activator activity | Cilium biogenesis/degradation<br>Endocytosis<br>Protein transport<br>Transport | -0.76 | 0.009 |
| ENSECAG00000018683 | PCF11 | PCF11 cleavage and polyadenylation factor subunit | mRNA processing | mRNA 3'-end processing<br>mRNA cleavage<br>mRNA polyadenylation<br>mRNA splicing, via spliceosome<br>termination of RNA polymerase II transcription | -0.78 | 0.010 |
| ENSECAG00000012328 | STAT5A | signal transducer and activator of transcription 5A | Activator<br>DNA-binding | Transcription<br>Transcription regulation | -0.89 | 0.001 |
| ENSECAG00000014662 | SLX4 | SLX4 structure-specific endonuclease subunit | DNA damage<br>DNA recombination<br>DNA repair | DNA repair | -0.90 | 0.020 |
| ENSECAG00000028573 | TRIOBP | TRIO and F-actin binding protein | Actin-binding | Cell cycle<br>Cell division<br>Mitosis | -0.99 | 0.014 |
| ENSECAG00000016843 | SETD5 | SET domain containing 5 | Chromatin regulator<br>Methyltransferase<br>Transferase | Transcription<br>Transcription regulation<br>Histone H3-K36 trimethylation<br>Regulation of histone acetylation<br>Regulation of chromatin organization | -1.02 | 0.023 |
| ENSECAG00000008726 | NPY | neuropeptide Y | Neuropeptide<br>Calcium channel regulator<br>activity<br>G protein-coupled receptor<br>activity<br>Hormone activity<br>signaling receptor binding | Adult feeding behavior<br>G protein-coupled receptor signaling<br>pathway<br>Innate immune response<br>Central nervous system neuron<br>development | -1.27 | 0.001 |
| ENSECAG00000021312 | ABHD14A | abhydrolase domain<br>containing 14A | Hydrolase |  | -1.30 | 0.007 |
| ENSECAG00000030141 | PXMP2 | peroxisomal membrane<br>protein 2 | Transporter | Protein import into peroxisome<br>membrane | -1.37 | 0.019 |
| ENSECAG00000014057 | ASB1 | ankyrin repeat and SOCS box<br>containing 1 | Developmental protein | Ubiquitine-like conjugation pathway | -1.42 | 0.006 |
| ENSECAG00000022130 | CPT2 | carnitine palmitoyltransferase<br>2 | Acyltransferase | Fatty acid metabolism<br>Lipid metabolism<br>Transport | -1.52 | 0.004 |
| ENSECAG00000015074 | LIPE | lipase E, hormone sensitive<br>type | Hydrolase | Cholesterol metabolism<br>Lipid degradation<br>Lipid metabolism<br>Steroid metabolism<br>Sterol metabolism | -1.53 | 0.018 |
| ENSECAG00000008600 | APOB | apolipoprotein B | Heparin-binding | Cholesterol metabolism<br>Lipid metabolism<br>Lipid transport<br>Steroid metabolism<br>Sterol metabolism<br>Transport | -1.99 | 0.041 |
| ENSECAG00000011240 | CNTN1 | Contactin 1 | Carbohydrate binding | Cell adhesion<br>Notch signaling pathway | -2.36 | 0.001 |

Log2 Fold-Change > 0 indicates up-regulation of the gene in embryos from old nulliparous mares; Log2 Fold-Change < 0 indicates down-regulation of the gene in embryos from old nulliparous mares

### Differential gene expression in the TE part

In the TE, 13,322 genes were considered as expressed in OM or ON. Twenty-three were differentially expressed (Additional file 2) with 16 genes being downregulated and 7 being up regulated in ON (Figure 2). Respectively, 14 and 6 out of the down- and upregulated genes were associated to a known protein in human. Moreover, despite the filtering, 2 down-regulated genes in ON (LIM and cysteine rich domains 1, *LMCD1*; lysophosphatidic acid receptor 4, *LPAR4*) were only expressed in one embryo and were not considered for further analysis. The remaining 19 genes are presented in Table 4. Down-regulated genes in the TE of old nulliparous mares were mainly involved in spindle organization, chromatin remodeling and cellular process, transcription and DNA repair, cell polarity, adhesions, junctions and signaling, extracellular matrix organization, ion homeostasis, glycerol metabolism, glycolysis, immunity, gastrulation and placenta development while up-regulated genes were related to ion binding, cell death, lipid metabolism, protein maturation and membrane organization

**Table 4:**
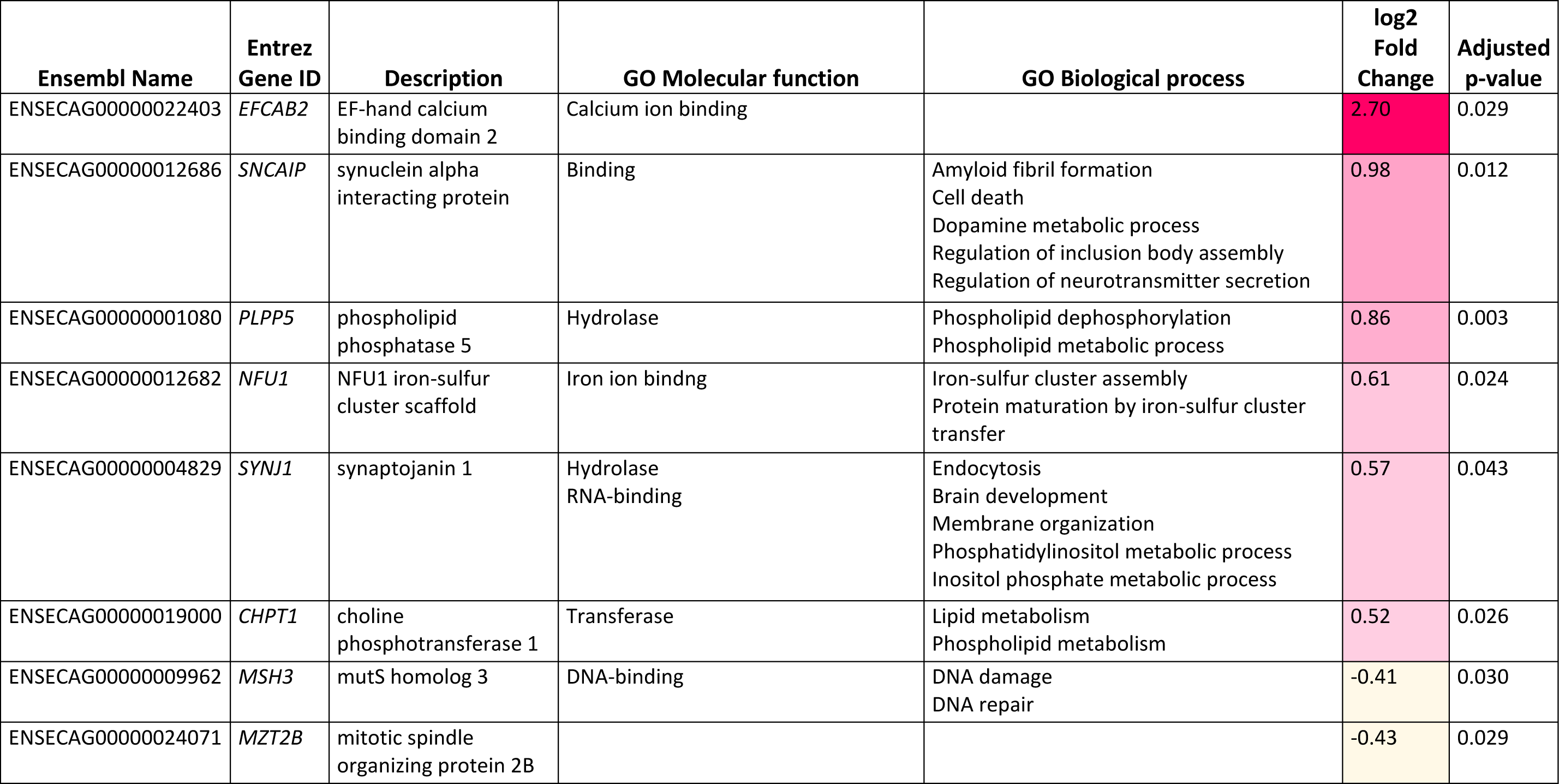
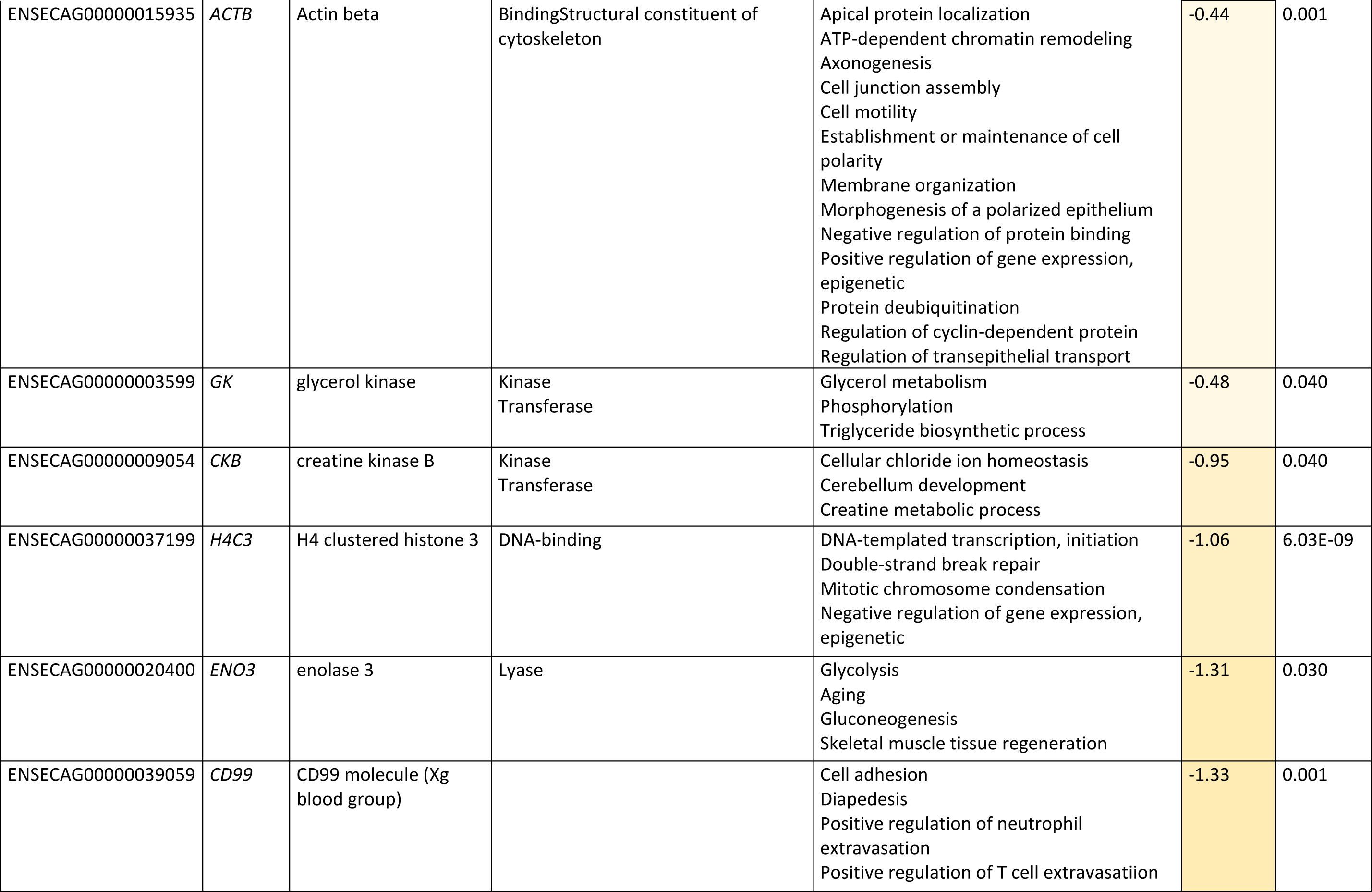
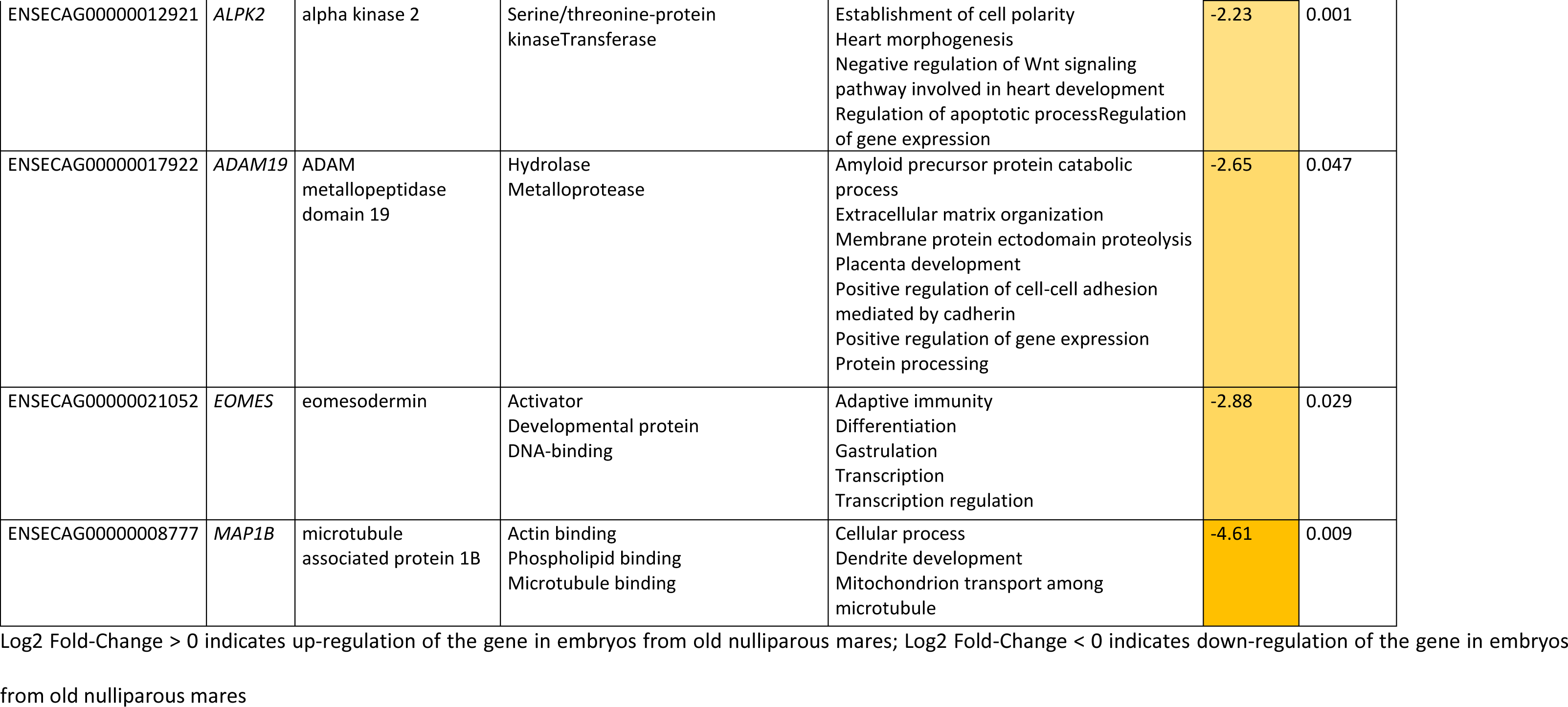
Up- and down-regulated genes coding for a protein in the trophoblast part of equine embryos acc ording to old mare parity

### Gene set enrichment analysis in deconvoluted ICM cells

After Entrez Gene ID conversion, 12,287 genes were considered expressed in ICM cells. Only one GO Biological Process and one KEGG pathways were disturbed by maternal parity in ICM cells (Additional file 3 & Table 5). The GO BP “Cytoplasmic microtubule organization” was enriched in the ICM cells from OM embryos (NES = -2.17). The KEGG pathway “Neuroactive ligand receptor interaction” was enriched in ICM cells from ON embryos (NES = 1.98). Detailed examination indicated that genes involved in this enrichment were related to biological regulation, signaling and response.

**Table 5:**
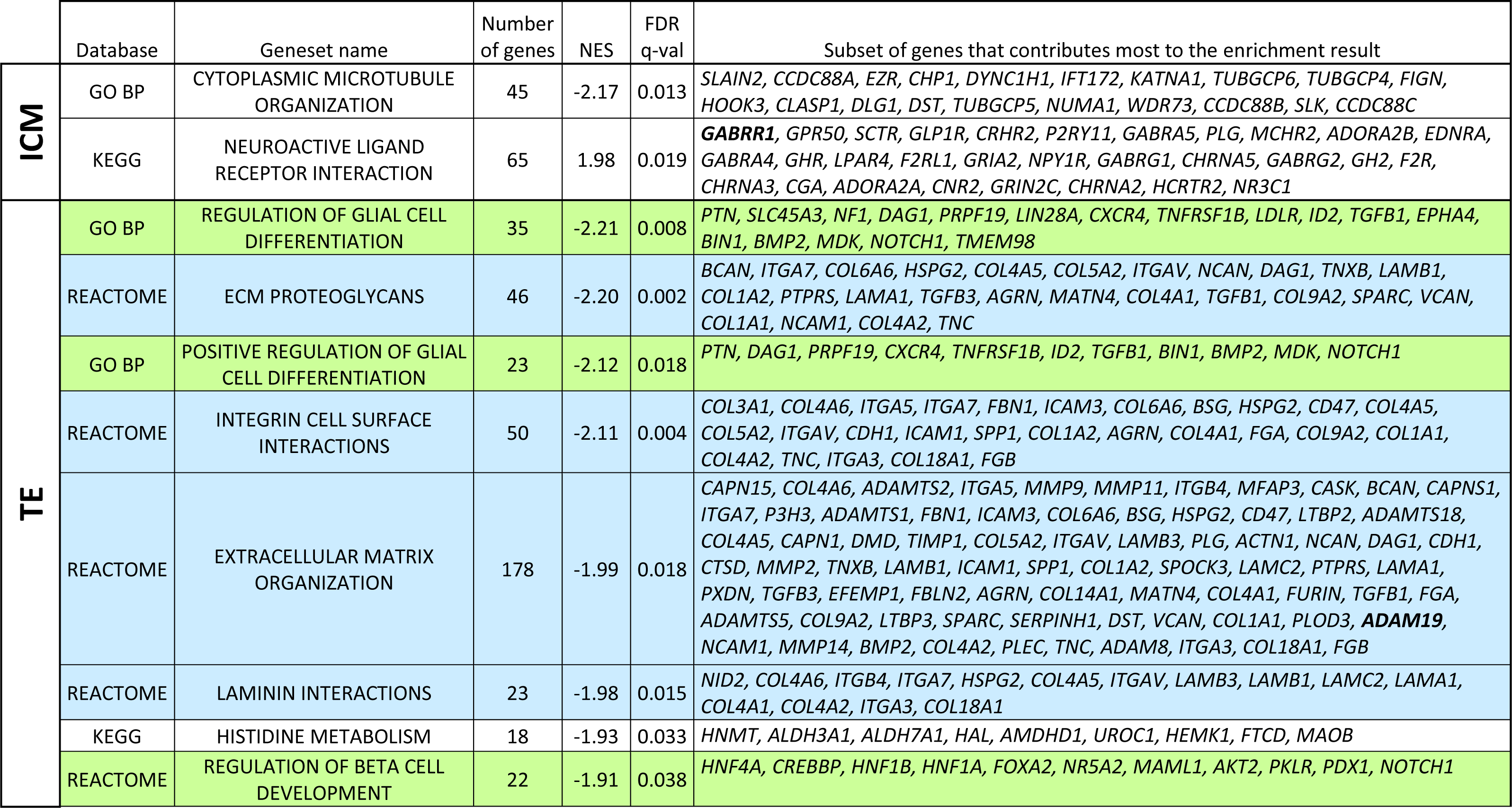
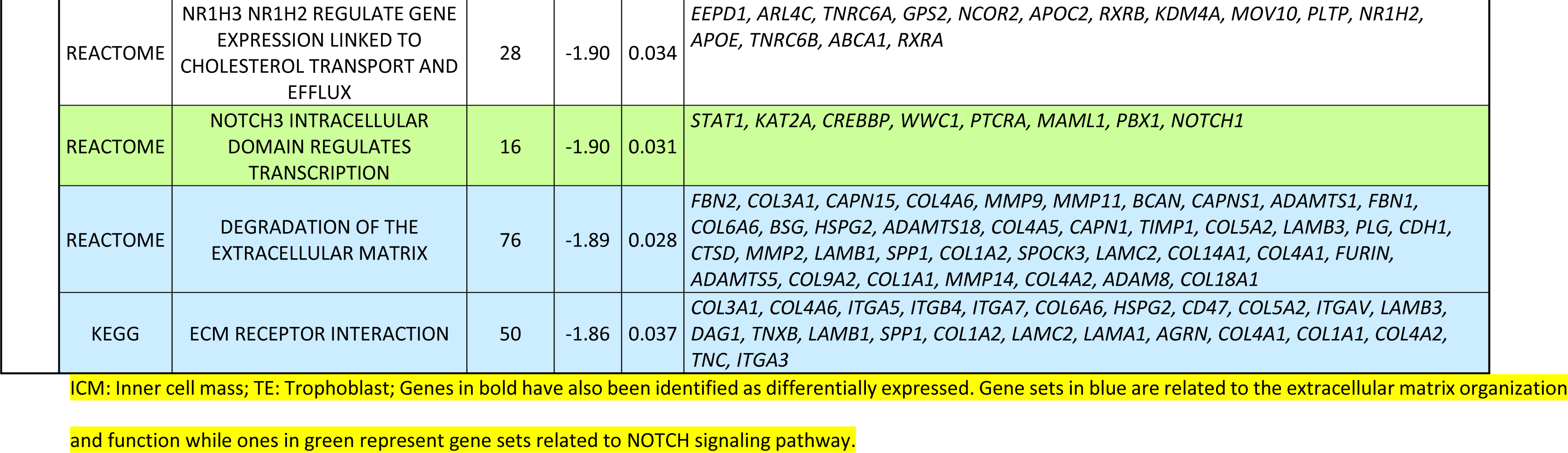
Enriched pathway in ICM and TE of embryos according to maternal parity in old mares

|  | Database | Geneset name | Number of genes | NES | FDR q-val | Subset of genes that contributes most to the enrichment result |
| --- | --- | --- | --- | --- | --- | --- |
| ICM | GO BP | CYTOPLASMIC MICROTUBULE ORGANIZATION | 45 | -2.17 | 0.013 | *SLAIN2, CCDC88A, EZR, CHP1, DYNC1H1, IFT172, KATNA1, TUBGCP6, TUBGCP4, FIGN, HOOK3, CLASP1, DLG1, DST, TUBGCP5, NUMA1, WDR73, CCDC88B, SLK, CCDC88C* |
|  | KEGG | NEUROACTIVE LIGAND RECEPTOR INTERACTION | 65 | 1.98 | 0.019 | *GABRR1, GPR50, SCTR, GLP1R, CRHR2, P2RY11, GABRA5, PLG, MCHR2, ADORA2B, EDNRA, GABRA4, GHR, LPAR4, F2RL1, GRIA2, NPY1R, GABRG1, CHRNA5, GABRG2, GH2, F2R, CHRNA3, CGA, ADORA2A, CNR2, GRIN2C, CHRNA2, HCRTR2, NR3C1* |
| TE | GO BP | REGULATION OF GLIAL CELL DIFFERENTIATION | 35 | -2.21 | 0.008 | *PTN, SLC45A3, NF1, DAG1, PRPF19, LIN28A, CXCR4, TNFRSF1B, LDLR, ID2, TGFB1, EPHA4, BIN1, BMP2, MDK, NOTCH1, TMEM98* |
|  | REACTOME | ECM PROTEOGLYCAN | 46 | -2.20 | 0.002 | *BCAN, ITGA7, COL6A6, HSPG2, COL4A5, COL5A2, ITGAV, NCAN, DAG1, TNXB, LAMB1, COL1A2, PTPRS, LAMA1, TGFB3, AGRN, MATN4, COL4A1, TGFB1, COL9A2, SPARC, VCAN, COL1A1, NCAM1, COL4A2, TNC* |
|  | GO BP | POSITIVE REGULATION OF GLIAL CELL DIFFERENTIATION | 23 | -2.12 | 0.018 | *PTN, DAG1, PRPF19, CXCR4, TNFRSF1B, ID2, TGFB1, BIN1, BMP2, MDK, NOTCH1* |
|  | REACTOME | INTEGRIN CELL SURFACE INTERACTIONS | 50 | -2.11 | 0.004 | *COL3A1, COL4A6, ITGA5, ITGA7, FBN1, ICAM3, COL6A6, BSG, HSPG2, CD47, COL4A5, COL5A2, ITGAV, CDH1, ICAM1, SPP1, COL1A2, AGRN, COL4A1, FGA, COL9A2, COL1A1, COL4A2, TNC, ITGA3, COL18A1, FGB* |
|  | REACTOME | EXTRACELLULAR MATRIX ORGANIZATION | 178 | -1.99 | 0.018 | *CAPN15, COL4A6, ADAMTS2, ITGA5, MMP9, MMP11, ITGB4, MFAP3, CASK, BCAN, CAPNS1, ITGA7, P3H3, ADAMTS1, FBN1, ICAM3, COL6A6, BSG, HSPG2, CD47, LTBP2, ADAMTS18, COL4A5, CAPN1, DMD, TIMP1, COL5A2, ITGAV, LAMB3, PLG, ACTN1, NCAN, DAG1, CDH1, CTSD, MMP2, TNXB, LAMB1, ICAM1, SPP1, COL1A2, SPOCK3, LAMC2, PTPRS, LAMA1, PXDN, TGFB3, EFEMP1, FBLN2, AGRN, COL14A1, MATN4, COL4A1, FURIN, TGFB1, FGA, ADAMTS5, COL9A2, LTBP3, SPARC, SERPINH1, DST, VCAN, COL1A1, PLOD3, **ADAM19**, NCAM1, MMP14, BMP2, COL4A2, PLEC, TNC, ADAM8, ITGA3, COL18A1, FGB* |
|  | REACTOME | LAMININ INTERACTIONS | 23 | -1.98 | 0.015 | *NID2, COL4A6, ITGB4, ITGA7, HSPG2, COL4A5, ITGAV, LAMB3, LAMB1, LAMC2, LAMA1, COL4A1, COL4A2, ITGA3, COL18A1* |
|  | KEGG | HISTIDINE METABOLISM | 18 | -1.93 | 0.033 | *HNMT, ALDH3A1, ALDH7A1, HAL, AMDHD1, UROC1, HEMK1, FTCD, MAOB* |
|  | REACTOME | REGULATION OF BETA CELL DEVELOPMENT | 22 | -1.91 | 0.038 | *HNF4A, CREBBP, HNF1B, HNF1A, FOXA2, NR5A2, MAML1, AKT2, PKLR, PDX1, NOTCH1* |

|  |  |  |  |  |  |
| --- | --- | --- | --- | --- | --- |
| REACTOME | NR1H3 NR1H2 REGULATE GENE<br>EXPRESSION LINKED TO<br>CHOLESTEROL TRANSPORT AND<br>EFFLUX | 28 | -1.90 | 0.034 | *EEPD1, ARL4C, TNRC6A, GPS2, NCOR2, APOC2, RXRB, KDM4A, MOV10, PLTP, NR1H2, APOE, TNRC6B, ABCA1, RXRA* |
| REACTOME | NOTCH3 INTRACELLULAR<br>DOMAIN REGULATES<br>TRANSCRIPTION | 16 | -1.90 | 0.031 | *STAT1, KAT2A, CREBBP, WWC1, PTCRA, MAML1, PBX1, NOTCH1* |
| REACTOME | DEGRADATION OF THE<br>EXTRACELLULAR MATRIX | 76 | -1.89 | 0.028 | *FBN2, COL3A1, CAPN15, COL4A6, MMP9, MMP11, BCAN, CAPNS1, ADAMTS1, FBN1, COL6A6, BSG, HSPG2, ADAMTS18, COL4A5, CAPN1, TIMP1, COL5A2, LAMB3, PLG, CDH1, CTSD, MMP2, LAMB1, SPP1, COL1A2, SPOCK3, LAMC2, COL14A1, COL4A1, FURIN, ADAMTS5, COL9A2, COL1A1, MMP14, COL4A2, ADAM8, COL18A1* |
| KEGG | ECM RECEPTOR INTERACTION | 50 | -1.86 | 0.037 | *COL3A1, COL4A6, ITGA5, ITGB4, ITGA7, COL6A6, HSPG2, CD47, COL5A2, ITGAV, LAMB3, DAG1, TNXB, LAMB1, SPP1, COL1A2, LAMC2, LAMA1, AGRN, COL4A1, COL1A1, COL4A2, TNC, ITGA3* |

ICM: Inner cell mass; TE: Trophoblast; Genes in bold have also been identified as differentially expressed. Gene sets in blue are related to the extracellular matrix organization
and function while ones in green represent gene sets related to NOTCH signaling pathway.

### Gene set enrichment analysis in TE

After Entrez Gene ID conversion, 11,993 genes were considered expressed in TE from ON or OM embryos. All the 2, 2 and 8 perturbed gene sets from GO BP, KEGG and REACTOME, respectively, were enriched in the TE of embryos from OM mares (NES < -1.8; Additional file 4 & Table 5). In the TE, several gene sets were related to extracellular matrix organization and function. Others were involved in NOTCH signaling pathway. The last two pathways were involved in histidine metabolism and cholesterol transport.

## Discussion

Maternal parity in mares older than 10 years old did not influence embryo recovery rates. This rate, however, seems lower in old nulliparous compared to old multiparous mares. The effect of mare nulliparity on the fertility in mares older than 10 years old have never been explored. For 3 mares, a developing embryo was observed at 14 days post ovulation, demonstrating that embryos were not recovered with the flushing. This happened only in multiparous mares and increased occurrence in multiparous mares has not been described to the author’s knowledge yet. Uterine size as well as histological and morphological degenerations in uterus could explain this result. After foaling, uterine size decreases quickly during the first week post-partum but the complete involution of both uterine horns seems to end only after twenty to thirty days [62,63] whereas histology of the uterine body endometrium returns to normal by 7 to 10 days post-partum [31]. Ageing and the multiplication of foalings, with repetition of uterine extensions and involutions may affect the uterus’s ability to involute. Indeed, clinically the authors have observed that a larger volume of fluid is required to flush the uterus of multiparous mares compared to young mares. Histologically, however, parity is correlated with the presence of elastosis in the myometrial vessels [37]. These alterations may be associated to alterations of uterine contractility and decrease fluid clearance in multiparous mares, both factors that could affect embryo recovery.

In both ICM and TE, only a few genes were affected by maternal parity. Some downregulated genes in embryos collected from ON mares were involved in common functions between ICM and TE such as regulation of transcription, cell cycle and development, cell organization and immunity. Up-regulated genes in the ICM of ON embryos were related to ion transport, regulation of transcription and catabolism of lipids, proteins and carbohydrates. In the TE, downregulated genes in ON embryos were related to glycerol metabolism, glycolysis and directly involved in gastrulation and placental formation while upregulated genes were related to cell death, lipid metabolism, protein maturation, membrane organization and as in ICM, ion binding.

In the GSEA analysis, only two gene sets were perturbed in the ICM in relation to maternal parity in old mares with one related to microtubule organization being enriched in ON and the other one being enriched in OM embryos and related membrane receptors. No gene set was enriched in the TE of ON mares but enriched pathways in OM embryos were mainly related to extracellular matrix organization and cell differentiation, mainly related to NOCTH signaling pathway.

Of particular interest, the gene SLX4 Structure-Specific Endonuclease Subunit (*SLX4*), the regulatory subunit of structure-specific endonucleases that are required for repair of DNA lesions, is down-regulated in the ICM of embryos from ON mares. In TE, moreover, part of the post-replicative DNA mismatch repair system, MutS Homolog 3 (*MSH3*) is downregulated in embryos from ON mares. These results indicate that DNA repair systems in both ICM and TE are affected by maternal parity. Interestingly, the gene Gamma-aminobutyric acid type A receptor subunit rho1, *GABRR1*, was not expressed in embryos collected from multiparous mares while in nulliparous mares, 4/5 embryos expressed this gene in the ICM. *GABRR1* encodes for a Cl^−^ channel receptor involved in the gamma-aminobutyric acid (GABA) pathway. Work on mouse embryonic and peripheral neural crest stem cells have shown that GABA receptors negatively affect preimplantation embryonic growth by negatively controlling cell proliferation, being involved in DNA damage checkpoint and by increasing cellular arrest in the S phase [64]. Alterations in the embryo environment because of mare parity could therefore modify DNA lesion repair and therefore, cell proliferation, suggesting that embryo growth is reduced in nulliparous mares.

Several gene sets related to NOTCH signaling pathways were also enriched in the TE of OM embryos and Notch Receptor 1 (*NOTCH1*) always contributed to those enrichments. NOTCH signaling pathway is essential for proper development, with *NOTCH1*being required for cell proliferation in early bovine embryos [65]. The enrichment of this pathway in embryos from multiparous mares therefore suggests that cell proliferation is slowed down in embryos from nulliparous mares

The expression of SET domain containing 5 (*SETD5*) signal transducer and activator of transcription 5A (*STAT5A*), PCF11 cleavage and polyadenylation factor subunit (*PCF11*), ribosomal protein L36 (*RPL36*), exosome component 7 (*EXOSC7*), H2A clustered histone 20 (*H2AC20*) were downregulated in the ICM and H4 clustered histone 3 (H4C3) were downregulated in the TE of embryos collected from nulliparous mares. These genes are all involved in transcription and/or translation, suggesting that the expression of genes in embryos from nulliparous mares was altered. Of particular interest, *SETD5* et STAT5A are known to be, respectively, key regulators of methylation and signaling via cytokines, both gene expressions being essential for embryo development [77,78].

In the TE, the gene encoding for eomesodermin (*EOMES*) is downregulated in embryos collected from ON mares. This gene controls the formation of germ cell layers and is involved in the differentiation of the trophoblast in the mouse [66] while in human, cattle and pigs, *EOMES* is not expressed in the preimplantation embryos [67-69]. In horses, *EOMES* is suggested to also be a marker of induced trophoblast cells but its role has never been explored [70]. If its role is similar as in mouse, this down-regulation could lead to a reduced differentiation of cells in the trophoblast, that could impair its principal function, i.e., the regulation of exchanges with the maternal environment.

As the external part of the embryo is exclusively composed of trophoblast in the mare at the studied developmental stage, poorer maternal-embryo exchanges through the trophoblast in nulliparous mares could explain the defects observed in both compartments. Indeed, the reduced gene expression of TRIO and F-Actin Binding Protein (*TRIOBP*) in the ICM as well as actin beta (*ACTB*) and microtubule associated protein 1B (*MAP1B*) in the TE of equine embryos seems to fit the hypothesis that molecule transfer is altered in embryos from ON mares. In polarized epithelial cells, such as the trophoblast, cytoskeleton is essential for the communication with the extracellular environment (for review [71]). The protein encoded by *ACTB* is a direct component of the cytoskeleton and the one encoded by *MAP1B* is a molecule responsible for the stabilization of microtubules [72]. *TRIOBP* regulates actin cytoskeletal organization and the formation of a Tara and TRIO complex coordinates actin remodeling which is essential for exchanges [73]. In addition, the extracellular matrix (ECM) is very important for embryo development and embryo-maternal exchanges (for a review [74]). ECM relative gene sets appeared to be altered by maternal parity in old mares with several gene sets enriched in the TE of embryos from ON mares. The expression of the ADAM metallopeptidase domain 19 gene (*ADAM19*), moreover, was reduced in the TE of ON embryos. This gene encodes for a transmembrane glycoprotein that is essential for tight junction formation. Tight junctions formation and integrity are essential for blastocyst development in mouse and pigs [75]. The reduction of the expression of *ADAM19* in the TE of embryos from old nulliparous mares, could therefore support the hypothesis that embryo integrity is altered, leading to alteration of ion and nutrient exchanges in both TE and ICM. Alterations of cytoskeleton, ECM and integrity of the TE in ON embryos, probably related to adaptation to the embryo environment, could affect embryo-maternal exchanges and consequently embryonic development.

The lipid metabolism is particularly important for mammalian embryo development as it is an important source of energy for growth (for review [79]). In horses, the embryo is particularly reliant on its environment as it develops free inside the mare’s uterus until around 35 days post ovulation, when it finally starts to implant. Several genes related to lipid metabolism and transport were affected in the ICM by maternal parity. Among them, carnitine palmitoyltransferase 2 (*CPT2*) is downregulated in the ICM of ON embryos. The protein encoded by this gene catalyzes the oxidation of fatty acids in the mitochondria. In mouse embryos, *CTP2* transcription is essential and could be used as a marker for future implantation [80]. Therefore, the reduced expression of this gene could indicate that embryo development is reduced because of reduced lipid oxidation in old nulliparous mares. This could be the result or at least related to reduced lipolysis via lipase or a defect in the transport of lipids. Both functions are altered in the ICM of ON embryos. Indeed, lipase E (*LIPE*), also known as hormone-sensitive lipase, and apolipoprotein B (*APOB*), involved in the transport of lipids, are downregulated in the ICM of ON embryos. APOB has been identified as very important for equine embryo development as its expression is increased by a factor of 200 between day 8 and day 14 [81]. Furthermore, the production of the protein apolipoprotein B by the endoderm is required for the development of preimplantation mouse embryos [82]. In the equine species, *APOB* expression is dysregulated in Day 8 embryos when mares do not produce enough progesterone [83]. It has been observed that the risk of embryo mortality is increased in mares producing less than 4 ng/ml of progesterone through the post ovulation period [84]. Progesterone, indeed, is mandatory for embryo development by modulating its environment to ensure pregnancy [85]. Reduced expression of *APOB* suggests that the expression of this gene is particularly dependent on the embryo environment. Here, except one, close to the cut-off value, all mares produced sufficient progesterone for normal gestation at 8 days post ovulation and there was no difference in progesterone concentration according to mare parity. It could be hypothesized, however, that the environment in the uterus of old nulliparous mares could lead to the difference of expression of this gene, independently of progesterone production.

## Conclusion

Mare’s parity, especially in mares older than 10 years old, has never been considered for the study of embryos. Here, however, it has been shown that mare’s parity in old mares affects the expression of genes in ICM and TE of blastocysts. Although only the expression of a few genes is altered by mare’s parity, some of these genes are particularly important for embryo growth and development. Genes related to nutrient exchanges and responses to environment signaling in both ICM and TE are particularly affected by the nulliparity of mares, suggesting that the developing environment from these mares are not optimal for embryonic growth. It is not possible to conclude, however, if differences in oocyte quality also play a role in those observations. More work on the oocyte, oviductal and uterine environment are needed to elucidate the exact mechanism leading to the present observations in Day 8 embryos.

In the present experiment, the destruction of the embryos made it impossible to predict their individual chance of implantation. The observed alterations suggest that implantation defects may be present in the embryos of old nulliparous mares. It is often assumed that nulliparous mares have a better uterine environment, more favorable to the development and implantation of an embryo than multiparous mares. In this study, however, being nulliparous and old does not seem to be the perfect match for embryonic development. If embryos succeed to implant, the apparent lower quality of embryos in nulliparous mares may accentuate differences in placentation and development observed in nulliparous mares, exacerbating the observed phenotype of smaller and lighter foals at birth.

## Supporting information

In manuscript

## Acknowledgments

The authors are grateful to the staff of the Institut Français du Cheval et de l’Equitation (IFCE) experimental farm (Jumenterie du Haras du Pin, Exmes, France) for care and management of animals. We acknowledge the high-throughput sequencing facility of I2BC for its sequencing and bioinformatics expertise. The bioinformatics analyses were performed thanks to Core Cluster of the Institut Français de Bioinformatique (IFB) (ANR-11-INBS-0013). The authors thank A.L. Lainé and the staff of the Phenotyping-Endocrinology laboratory for the hormonal assays. Many thanks to Matthias Zytnicki and Christophe Klopp for their advice on RNA-seq de novo analysis. Many thanks to Pablo Ross who kindly provided the coordinates for the XIST gene.

## Author contributions

PCP obtained the funding. PCP and VD conceived the project. FDG, LB, VD and PCP supervised the study. ED, CG, AM, CA, ND, NP, FDG, VD and PCP adapted the methodology for the project. ED, CG, AM, CA and YJ performed the experiments. AM, CG, CA, ND, NP, MD, LB and FDG provided the resources. ED, LJ, YJ and RL performed data curation. ED and LJ analyzed the data. ED wrote the original draft. All authors read, revised, and approved the submitted manuscript.

## List of abbreviations

DEG: differential expressed genes
DeMixT_ICM_cells: deconvoluted gene expression in ICM cells
DeMixT_TE_cells: deconvoluted gene expression in TE cells
ECM: Extracellular matrix
ERR: embryo collection rate
FDR: false discovery rate
GO BP: Gene Ontology biological process
GO: Gene Ontology
GSEA: gene set enrichment analyses ICM: inner cell mass
ICMandTE: inner cell mass enriched hemi-embryo ICSI: intracytoplasmic sperm injection
KEGG: Kyoto Encyclopedia of Genes and Genomes Log2FC: log2 fold change
NES: normalized enrichment score
OM: old multiparous mares
ON: old nulliparous mares
TE: trophoblast
TE_part: pure trophoblast hemi-embryo
XIST: X inactive Specific Transcript

## Additional file 1: Supp1.xlsx

Differential gene analysis using DeSeq2 in DeMixT_ICM_cells of equine embryo at Day 8 post-ovulation according to mares parity

Equine ensemble ID, orthologue human Ensembl ID, Orthologue human Entrez Gene ID, gene description, normalized counts for each embryo and parameters obtained after Deseq2 analysis (log2FoldChange, pvalue and padj (after FDR correction)) of genes expressed in ICM (after gene expression deconvolution of ICMandTE using DeMixT) of ON and OM embryos

ICM: Inner cell mass; ON: Old nulliparous mares; OM: old multiparous mares

## Additional file 2: Supp2.xlsx

Differential gene analysis using DeSeq2 in TE_part of equine embryo at Day 8 post-ovulation according to mares parity

Equine ensemble ID, orthologue human Ensembl ID, Orthologue human Entrez Gene ID, gene description, normalized counts for each embryo and parameters obtained after Deseq2 analysis (log2FoldChange, pvalue and padj (after FDR correction)) of genes expressed in TE_part of ON and OM embryos

ICM: Inner cell mass; ON: Old nulliparous mares; OM: old multiparous mares

## Additional file 3: Supp3.xlsx

Gene set enrichment analysis results on gene expression of DeMixT_ICM_cell of embryos from old nulliparous and multiparous mares

Gene Set Enrichment Analysis results (database, pathway name, size, enrichment score without and with normalization, p-value and FDR corrected q-value) for GO biological process, KEGG and REACTOME databases on genes expressed in ICM (after gene expression deconvolution of ICMandTE using DeMixT).

ICM: Inner cell mass

## Additional file 4: Supp4.xlsx

Gene set enrichment analysis results on gene expression of TE_part of embryos from old nulliparous and multiparous mares

Gene Set Enrichment Analysis results (database, pathway name, size, enrichment score without and with normalization, p-value and FDR corrected q-value) for GO biological process, KEGG and REACTOME databases on genes expressed in TE_part

TE: trophoblast

